# Structure modeling and specificity of peptide-MHC class I interactions using geometric deep learning

**DOI:** 10.1101/2022.12.15.520566

**Authors:** Alon Aronson, Tanya Hochner, Tomer Cohen, Dina Schneidman-Duhovny

## Abstract

Major Histocompatibility Complex (MHC) plays a major role in the adaptive immune response by recognizing foreign proteins through binding to their peptides. In humans alone there are several hundred different MHC alleles, where each allele binds a specific subset of peptides. The peptide-MHC complex on a cell surface is identified by a T-cell receptor (TCR) and this binding invokes an immune response. Therefore, predicting the binding specificity of peptide-MHC pairs is necessary for understanding the immune recognition mechanism. Here, we develop an end-to-end novel deep learning model, MHCfold, that consists of structure and specificity prediction modules for simultaneous modeling of peptide-MHC class I (pMHCI) complexes and prediction of their specificity based on their modeled structure. MHCfold produces highly accurate structures of pMHCI complexes with mean Cα RMSD of 0.98Å and 1.50Å for the MHC α chain and the peptide, respectively. The binding specificity is also predicted with high accuracy (mean AUC of 0.94). Furthermore, the structure modeling component is orders of magnitudes faster than state-of-the-art methods (modeling of 100,000 pMHCI pairs in four hours on a standard computer), enabling high-throughput applications for large immunopeptidomics datasets. While peptide-MHC specificity can be accurately predicted from the sequence alone, TCR specificity prediction likely requires modeling of the 3D structures. We anticipate our model can be further used in structure-based prediction of TCR specificity.

MHCfold is available @https://github.com/dina-lab3D/MHCfold

## Introduction

Cell surface antigen peptide presentation by the major histocompatibility complex (MHC) is an important step in the adaptive immune response that is followed by a T cell receptor (TCR) recognition ^1^. There are two main MHC subgroups: class I proteins are expressed in all cells and present protein peptides of cytosolic and nuclear origin and class II proteins are expressed only in antigen representing cells (APC), where they bind exogenous antigenic peptides that originate extracellularly from pathogens. MHC class I (MHCI) has a closed shape binding groove on its α chain that can accommodate peptides of 8-10 amino acids. In contrast, the binding groove of MHC class II (MHCII) is open ended and spanned by both α and β chains, accommodating longer peptides of 11-20 amino acids. MHC proteins are polymorphic, and the various alleles, while similar in structure, have different peptide binding specificities ^2^. Moreover, the high diversity of TCRs, resulting from the V(D)J gene recombination, is estimated at ~10^15^ for T cells ^3^. This diversity enables our immune system to recognize epitopes from a near infinite number of antigens. Decoding the specificity between MHCs, peptides, and TCRs can have implications in cancer immunotherapy and in designing vaccines and biological therapies ^4^. Specifically, identification of neoantigens, that are peptides containing tumor-specific mutations, can aid in enhancing cancer immunotherapy ^5^.

Most of the available methods that predict peptide-MHC binding specificity rely on the sequences of MHC alleles and known peptide binders to train specificity predictors. The most commonly used method, NetMHCpan ^1,6^, trains an ensemble of dense neural networks with one hidden layer. Additional methods include MHCFlurry, MHCAttnNet, HLAthena and others that rely on Convolutional Neural Networks (CNNs) or Attention modules ^7–11^. While these methods are highly accurate (mean AUC of about 0.90-0.95 for MHCI peptides) mainly due to large datasets of known peptide-MHC pairs, they do not rely on structural information. To integrate structural data two bottlenecks need to be addressed. First, a highly accurate and high-throughput structure prediction protocol is necessary for modeling millions of peptide-MHC pairs. Second, structural information is significantly richer compared to sequences and requires more complex network architectures for machine learning. Structure-based prediction of specificity might further improve the accuracy of binding classification ^12,13^. In addition, accurately predicted peptide-MHC structures can be further used for predicting TCRs binding modes and specificity ^14^.

Recent progress in structure prediction resulted in novel highly accurate deep learning-based models for an end-to-end structure prediction, including AlphaFold2 ^15^ and RosettaFold ^16^. To achieve high accuracy, in addition to input sequence, evolutionary information based on a multiple sequence alignment (MSA) is used. However, this information is not available for establishing contacts between the highly variable peptides that can bind to the same MHC allele. Recently, a method that relies on AlphaFold2 to predict the peptide-MHC structure and classify binders and non-binders was released ^17^. The method uses available template structures from the Protein Data Bank (PDB) ^18^ without MSA to achieve high accuracy in structure modeling. Further, a logistic regression layer converts the complex predicted aligned error (PAE) values into binding/non-binding classification.

To produce 3D structures, both AlphaFold2 ^15^ and RosettaFold ^16^ use a specialized structure module that is invariant to transformations. However, for predicting structures of the same fold, such as MHC (Fig. 1A), we can achieve transformational invariance by aligning the structures of the training set ^19^. This enables structure prediction with a much simpler architecture and significantly lower train and inference times. Moreover, accurate models can be obtained without MSA.

**Figure 1.**
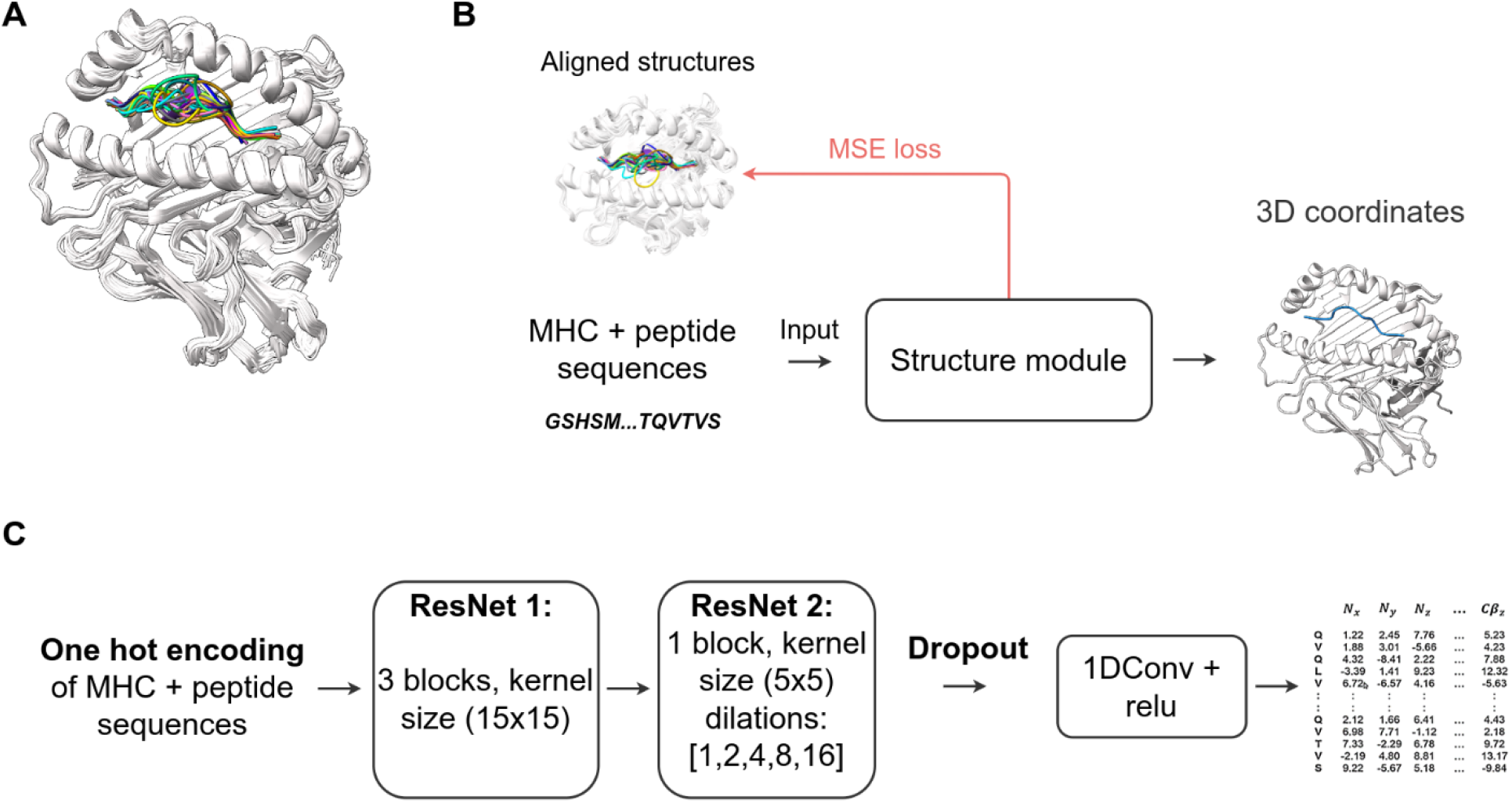
Structure module. **A.** The representative aligned pMHCI structures from the PDB. The aligned structures are used to achieve transformational invariance for the training of the model. **B.** MHCfold structure module: the input is the MHC and peptide sequences and the output is the 3D structure of their complex. **C.** Schematic architecture of MHCfold structure module.

Here, we design a dedicated network module with simple ResNet architecture for rapid and accurate prediction of pMHCI structures. We integrate the structure prediction and specificity classification modules for simultaneous modeling of pMHCI complexes and prediction of their specificity based on their modeled structure. We achieve high-accuracy structure predictions with a mean Cα RMSD of 0.98Å and 1.50Å for the MHC α chain and the peptide, respectively. The mean binding classification AUC is 0.94 on our test sets compared to 0.93 without the structural information.

## Results

### Structure prediction module

The input to the structure module are the sequences of α and chains of the MHC, and of a peptide, and the output is the coordinates of backbone and C atoms of the input sequences (Fig. 1B). The network was trained on a dataset of 390 pMHCI structures and validated on 41 complexes from the PDB (all solved prior to November 2021). The test set of 41 pMHCI structures was selected such that there is no peptide in the training set with more than 70% sequence identity for the given MHC allele. The structure between different MHC alleles is highly conserved with Cα RMSD under 1Å between aligned structures and a much higher variability in peptide conformations (Fig. 1A). Therefore, we achieved transformational invariance by aligning all the structures of the training set on an α chain of a randomly selected reference structure using the align tool from the Pymol viewer ^20^. This alignment enables the network to learn directly the pMHCI 3D coordinates in the coordinate frame of the reference structure. MHCfold is a convolutional neural network (CNN) that consists of two 1D Residual Neural Networks (ResNets)^21^ (Fig. 1C). The input sequences are represented using a one-hot encoding with 25 channels (Methods). The loss function is defined as a Mean Squared Error (MSE) on the coordinates of the backbone and Cβ atoms, which is equivalent to the squared RMSD. Once the coordinates of the backbone and Cβ atoms are predicted, the side chains are added using SCWRL ^22^ or MODELLER ^23^.

### MHCfold structure module produces high-accuracy models

We assess the accuracy of the predicted structures separately for the MHCI and the peptide because the MHCI structure is rigid and conserved while peptide conformations and their orientation in the binding groove show a significant variance (Fig. 1A). The predicted structures are superimposed on the experimental structure using the binding groove (first 177 amino acids of α chain) and the backbone RMSDs for the MHC and peptide are calculated. The binding groove is predicted with sub angstrom accuracy: the average Cα RMSD of the binding groove is 0.98Å and the median is 0.80Å on our test set. In contrast, the accuracy is lower for peptide conformations: the average Cα RMSD is 1.50Å and the median is 1.40Å (Fig. 2A,B).

**Figure 2.**
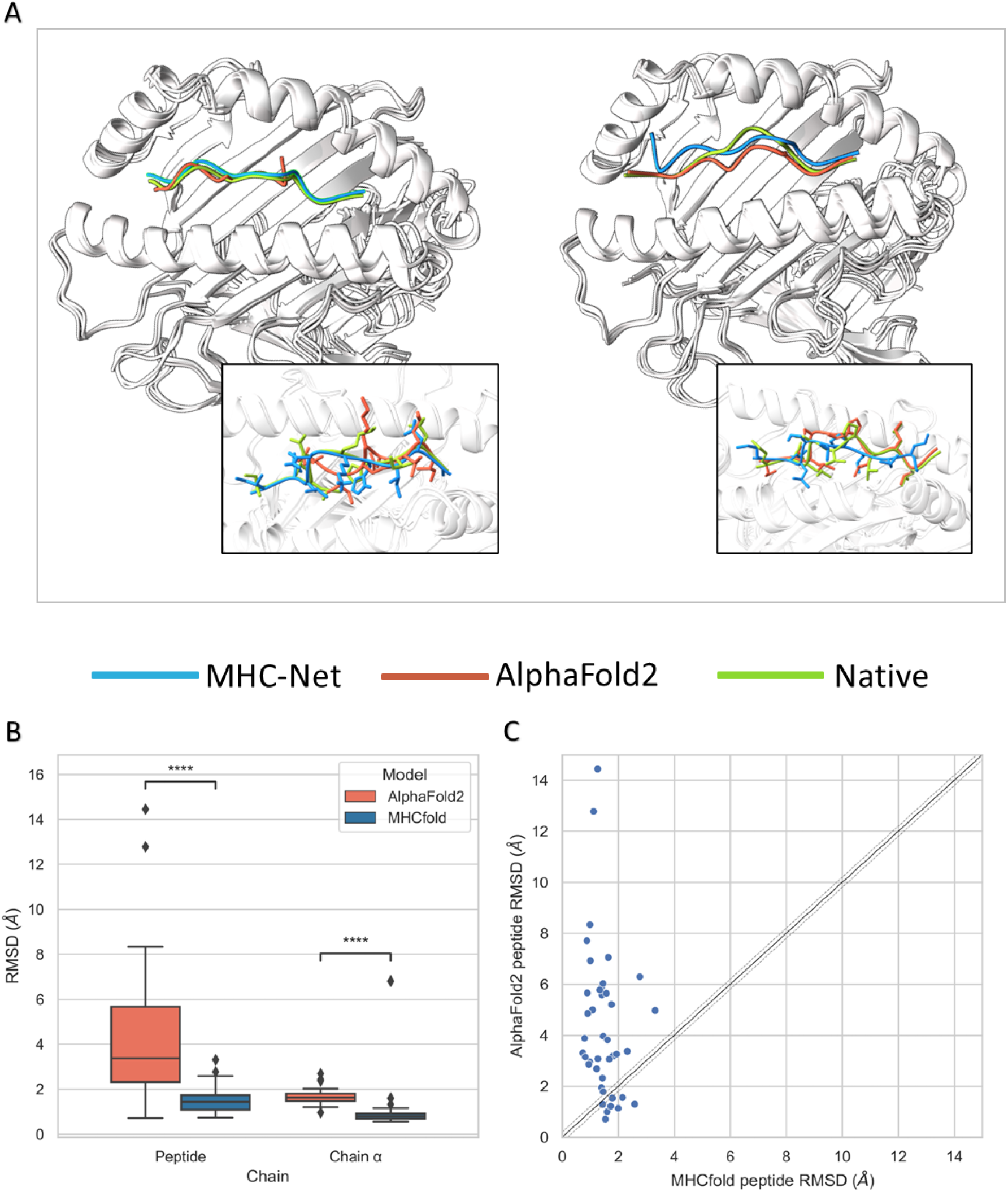
pMHCI structure modeling results. **A.** Two test set examples of modeled structures by MHCfold (blue), AlphaFold2 (orange) vs. experimental (green). PDB 7KGO (left): peptide RMSD 1.1Å for MHCfold vs. 12.8Å for AFM; 2CLR (right): peptide RMSD 2.5Å for MHCfold vs. 1.3Å for AFM. **B.** Boxplots of RMSDs of the MHC binding groove (residues 1-177 of the α chain) and of the peptides in the test set (41 pMHCI structures), chain α t-test p-value of 4.16e-5, peptide t-test p-value of 1.83e-6 **C.** Peptide RMSDs for MHCfold vs. AFM, each dot corresponds to a structure from the test set.

### Comparison to AlphaFold2

Recent developments in *ab initio* structure prediction ^15^ enable obtaining accurate structures also for protein-peptide complexes ^24,25^. We apply AlphaFold-Multimer (AFM) on our test set and compare the predicted structures to the ones obtained from MHCfold. Our test set contains 36 structures that were part of the AFM training set. AFM generated structures had a slightly higher α chain binding site RMSD (Fig. 2B, mean: 1.65Å, median: 1.62Å). Remarkably, the peptide RMSD for MHCfold was significantly lower than AFM (Fig. 2B-C, average: 1.50Å vs. 4.30Å, median: 1.40Å vs. 3.40Å for MHCfold and AFM, respectively). Moreover, there is a significant difference in the running times of both models. The average runtime of AFM for a single pMHCI structure prediction using the ColabFold version ^26^ in colab is ~150s. The average runtime of MHCfold for a single pMHCI structure prediction is ~ 0.15s without side-chain reconstruction. Side chain placement using SCWRL ^22^ takes around 1s. Overall, MHCfold is orders of magnitude faster than AFM for predicting pMHCI complexes.

### Specificity prediction module

Due to the majority of peptide specificity data originating from peptide elution experiments with only binary binding/non-binding labels, we formulate our problem as a classification. The input to the specificity prediction module is the sequences of the α and β chains of the MHC, and the peptide, and the output is the binary binding classification. The dataset used for this part was adapted from NetMHCpan 4.1 ^1^. It consists of 3,844,170 data points, divided into train and validation sets. The structure module is used as part of the classification module to produce a pairwise distance matrix of the predicted pMHCI complex. The classification module is based on a transformer’s encoder ^27^, with the additional distance information from the structure module injected at the multi-head attention module in the encoder.

### MHCfold binding module predicts with high accuracy

We evaluate the model using the AUC computed separately for each MHC allele in the test set. The mean AUC of the different alleles is 0.94, and the median is 0.95 (Table 1.). We performed an ablation study where we trained the MHCfold binding module without the pairwise distances injection, meaning the altered network’s architecture was that of a regular transformer. The performance without the structural information is slightly lower (Fig. 4A,B), indicating that the distances contribute to more accurate classification.

**Table 1.**
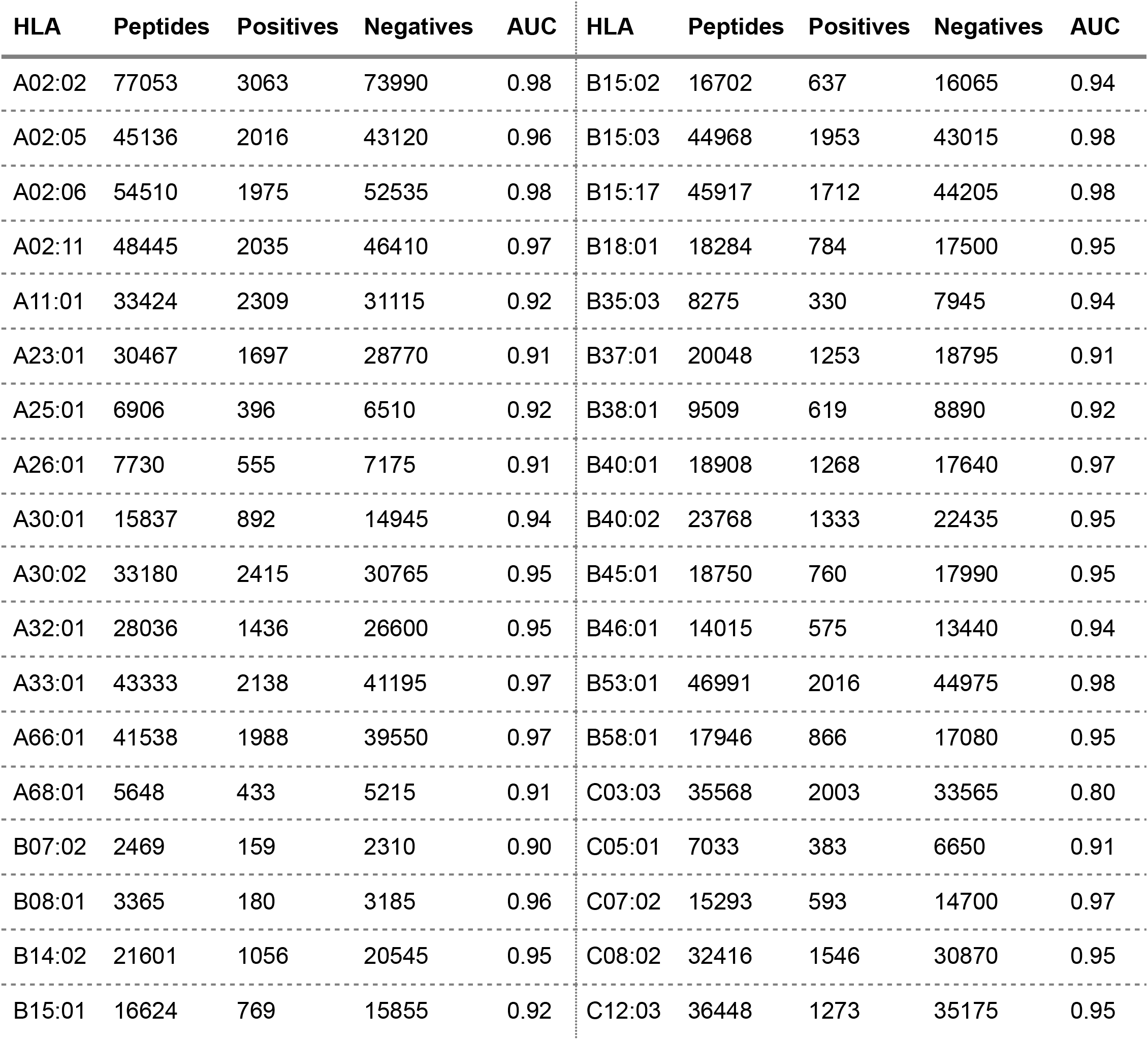
Performance of MHCfold for binding classification on the test set per MHC allele.

| HLA | Peptides | Positives | Negatives | AUC | HLA | Peptides | Positives | Negatives | AUC |
| --- | --- | --- | --- | --- | --- | --- | --- | --- | --- |
| A02:02 | 77053 | 3063 | 73990 | 0.98 | B15:02 | 16702 | 637 | 16065 | 0.94 |
| A02:05 | 45136 | 2016 | 43120 | 0.96 | B15:03 | 44968 | 1953 | 43015 | 0.98 |
| A02:06 | 54510 | 1975 | 52535 | 0.98 | B15:17 | 45917 | 1712 | 44205 | 0.98 |
| A02:11 | 48445 | 2035 | 46410 | 0.97 | B18:01 | 18284 | 784 | 17500 | 0.95 |
| A11:01 | 33424 | 2309 | 31115 | 0.92 | B35:03 | 8275 | 330 | 7945 | 0.94 |
| A23:01 | 30467 | 1697 | 28770 | 0.91 | B37:01 | 20048 | 1253 | 18795 | 0.91 |
| A25:01 | 6906 | 396 | 6510 | 0.92 | B38:01 | 9509 | 619 | 8890 | 0.92 |
| A26:01 | 7730 | 555 | 7175 | 0.91 | B40:01 | 18908 | 1268 | 17640 | 0.97 |
| A30:01 | 15837 | 892 | 14945 | 0.94 | B40:02 | 23768 | 1333 | 22435 | 0.95 |
| A30:02 | 33180 | 2415 | 30765 | 0.95 | B45:01 | 18750 | 760 | 17990 | 0.95 |
| A32:01 | 28036 | 1436 | 26600 | 0.95 | B46:01 | 14015 | 575 | 13440 | 0.94 |
| A33:01 | 43333 | 2138 | 41195 | 0.97 | B53:01 | 46991 | 2016 | 44975 | 0.98 |
| A66:01 | 41538 | 1988 | 39550 | 0.97 | B58:01 | 17946 | 866 | 17080 | 0.95 |
| A68:01 | 5648 | 433 | 5215 | 0.91 | C03:03 | 35568 | 2003 | 33565 | 0.80 |
| B07:02 | 2469 | 159 | 2310 | 0.90 | C05:01 | 7033 | 383 | 6650 | 0.91 |
| B08:01 | 3365 | 180 | 3185 | 0.96 | C07:02 | 15293 | 593 | 14700 | 0.97 |
| B14:02 | 21601 | 1056 | 20545 | 0.95 | C08:02 | 32416 | 1546 | 30870 | 0.95 |
| B15:01 | 16624 | 769 | 15855 | 0.92 | C12:03 | 36448 | 1273 | 35175 | 0.95 |

**Table 2.** MHCfold modules parameters.

|  | Structure model | Classification module |
| --- | --- | --- |
| Total parameters | 1,672,899 | 3,333,956 |
| Trainable parameters | 1,670,987 | 1,661,057 |
| Non -trainable parameters | 1,912 | 1,672,899 |

**Figure 3.**
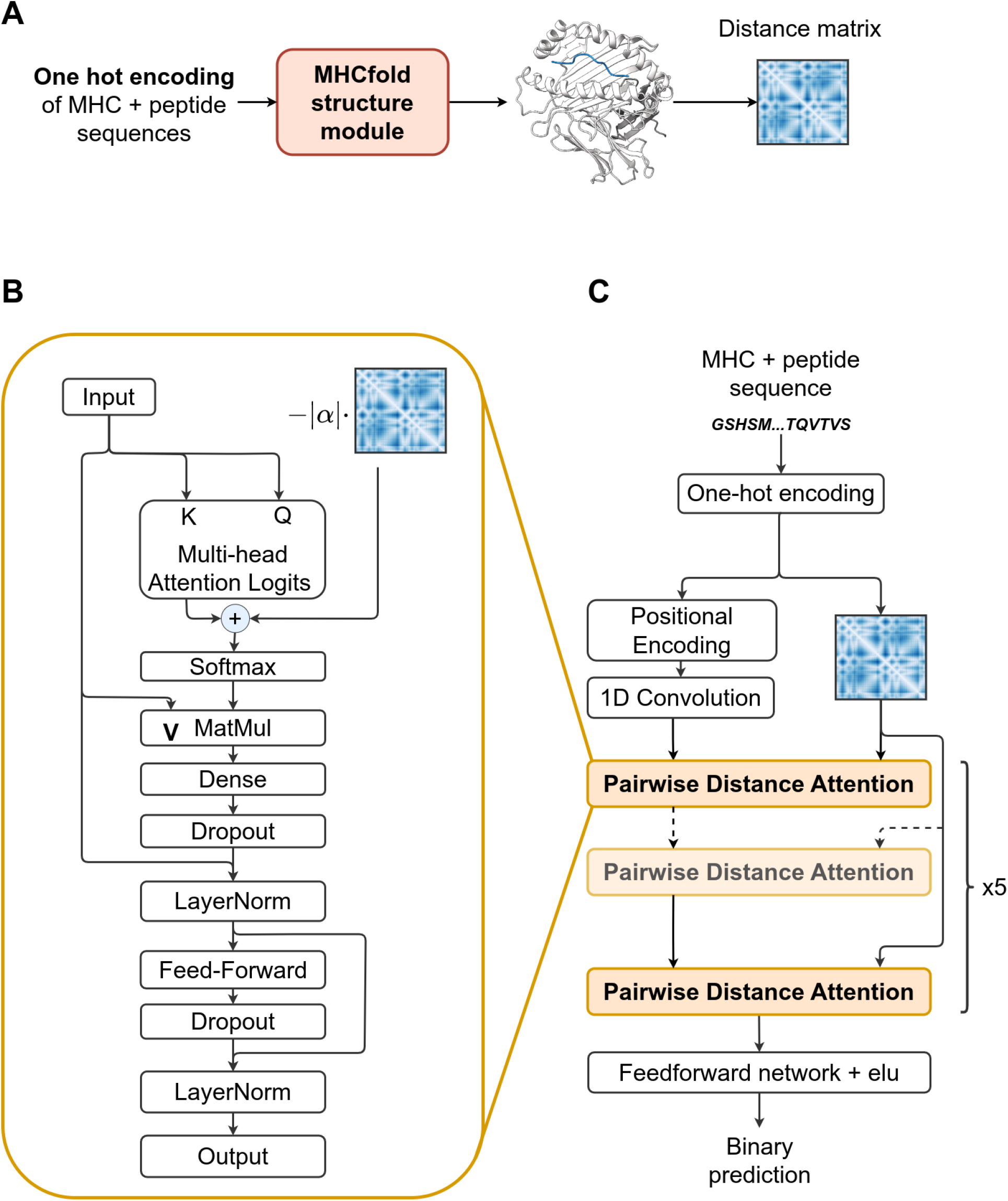
Specificity prediction module. **A.** The structure module is used to extract the pairwise distance matrix of the pMHCI. **B.** The pairwise distance attention component in the specificity prediction module. **C.** The architecture of the specificity prediction module. The input MHC and peptide sequences are represented by a one-hot encoding. The module is based on a transformer’s encoder, with additional distance information from the structure module injected at the multi-head attention module in the encoder. The output is the binary binding classification.

**Figure 4.**
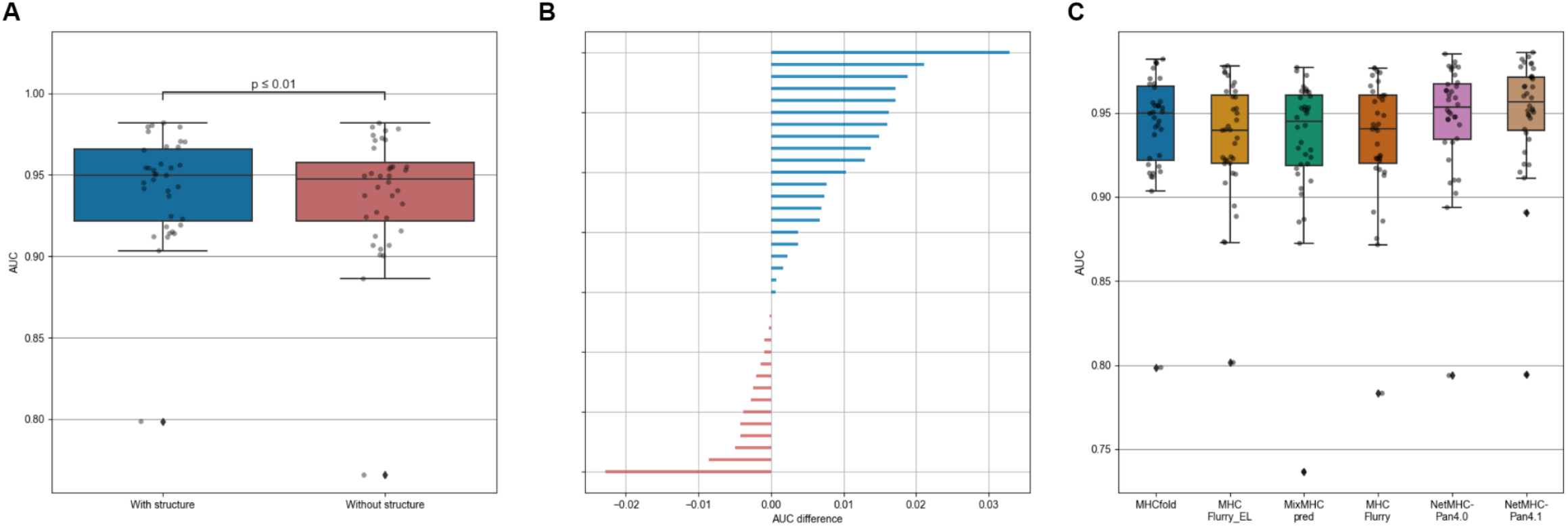
Specificity prediction performance. **A.** Comparison of the performance of the binding module with and without the structure module. Mean AUC with structure - 0.94, without - 0.93. **B.** The difference in AUC for each of the 36 test set alleles, between the MHCfold model with the structure component (positive difference, blue), and the model without (negative difference, red). **C.** Boxplots of AUC values for each MHC allele in the test set: MHCfold (blue), MHCFlurry_EL (yellow), MixMHCpred (green), MHCFlurry (orange), NetMHCpan4.0 (purple) and NetMHCpan4.1 (brown)

### Comparison to other classification methods

We compared MHCfold with state-of-the-art methods for binding specificity prediction that are based only on the sequences, including NetMHCpan 4.1 ^1^, MixMHCpred ^28^, and MHCFlurry ^29^. All the methods were tested on the same dataset. Overall, all the methods perform well with AUC above 0.93 (Fig. 4C). Most likely the high-performance is achieved due to a large number of available pMHCI pairs, especially from the EL mass spectrometry experiments.

## Discussion

We developed a high-throughput accurate end-to-end deep-learning based method for structure modeling and specificity prediction of pMHCI complexes. Integration of the structure generation module in the specificity prediction module enabled us to extract pairwise distances and use them as weights in a self-attention component, providing information in addition to sequences.

As the available structural data is limited, we used a simple architecture of ResNets, creating an end-to-end model that is trained on aligned structures. Compared to the state-of-the-art protein folding method AlphaFold2 ^15^, our structure module produces high accuracy models without MSA and (Fig. 2) with orders of magnitude lower runtimes, enabling high-throughput applications, including binding specificity prediction. We achieve low RMSD on the α chain (mean 0.98 Å), and higher RMSD on the peptide (mean 1.5Å), which is expected as most of the variability in solved pMHCI structures is in the peptide conformation.

This same approach is applicable to MHC class II modeling. As the MHC class II binds longer peptides, the model as it is is not suitable for that problem. We believe that with minor changes to the models architecture, it is possible to train a new model for peptide-MHC class II interactions, or maybe one model that can predict both classes.

We used our structure module to extract pairwise distances, and used them as weights in a self-attention component, so that the additional structure information would help the classification. Our results are objectively good, but in comparison to the state of the art NetMHCpan 4.1 ^1^, we still underperform a bit. We think this difference might be because we used only human data to train our model, where in their training they used data from other mammals as well. As an expected next step, we intend to train our model again with the data of other mammals, and expect an improvement in its performance.

In the current approach we relied on the straightforward one-hot representation of the sequence. This can be replaced by more sophisticated embedding that can be generated using sequence models, such as Restricted Boltzmann Machines or language models that enable improved predictions for rare alleles ^30,31^.

Our structural models of pMHCI enable the next step of modeling complexes with TCRs. We have previously used a similar approach for structure modeling of TCR Vβ domains ^19^. Integrating the two networks will enable modeling of the ternary peptide-MHC-TCR complexes, possibly with their specificities given a sufficient amount of training data. Our high-throughput approach enables rapid modeling of millions of complexes.

Our approach can also be used in the search for tumor-specific neoantigens, i.e., antigenic peptides carrying cancer-specific mutations ^32^. Prediction of such peptides from a large number of mutated protein sequences in the cancer cells can aid in identification of neoantigens and improve immunotherapy ^33,34^.

## Methods

### Structure module dataset

The pMHCI structures for training and testing this module were extracted from the PDB ^35^ with the release date prior to November 2021 and the resolution higher than 3.5Å. Duplicate structures and structures with peptide length shorter than 8 or longer that 10 were removed. The final dataset contains 431 experimentally solved structures of pMHCI complexes for various alleles of MHCI. We split the data into 90% for training and 10% for testing, using 70% peptide sequence identity cutoff, per allele, from the training set.

### Specificity prediction module dataset

The dataset used for training and testing the specificity prediction module was adapted from the recent NetMHCpan 4.1 dataset ^1^. The data was retrieved from either mass spectrometry (MS) experiments ^36–38^, known as eluted ligands (EL) or binding affinity (BA) data. The majority of the data is from the EL experiments with 3,679,405 pMHCI pairs and a small fraction of the data (170,470 pMHCI pairs) from the BA assays that include binding affinity. We binarized the BA data, using a threshold of 0.4256 similarly to the NetMHCpan 4.1 ^1^ to label the data points as positives (binders) and negatives (non-binders). After omitting duplicates in this dataset, we were left with 3,844,170 pMHCI pairs, which were divided into 90% for the training set and 10% for the validation set, with 3,459,753 and 384,417 data points respectively. The test set was also adapted from the NetMHCpan 4.1 ^1^, which is divided into 36 different MHCI alleles, with a varying amount of data points per allele (Table 1). The total number of data points in the test set is 946,141, where 45,416 are positively labeled, and 900,725 are negatively labeled.

### Sequence representation

The sequences are represented by an input tensor of 415×25, where 415 represents the maximal length of the concatenation of the three pMHCI complex chains. Each chain has a maximal length (α chain - 290, β chain - 110, and peptide - 15), and is padded with a special padding character if it is shorter than this length. The 25 channels are used as follows: 22 channels are used for the one-hot encoding of the 20 amino acids, including an unknown amino acid and the padding character. The three additional channels are used for indicating the chain (α, β, or peptide).

### Structure module architecture

This network module consists of two 1D ResNets ^21^ (Fig. S1). The first ResNet has a kernel of size 15, with 32 channels, and is fed into itself 3 times. Next we convert the tensors to 415×172 dimensions using a 1D convolution with 172 kernels. The second ResNet consists of a kernel of size 5 with dilated convolutions. We use five different dilation values (1, 2, 4, 8, and 16), and 172 channels. Finally, we convert the tensors to 415×64 and then to 415×15, using a convolution layer. This last tensor represents the output coordinates of the N, Cα, C, O and Cβ atoms and is compared directly to the actual coordinates to calculate loss. The whole network consists of 1,672,899 trainable parameters. The network was implemented using the TensorFlow library with keras^39,40^.

### Specificity prediction module architecture

The network for this module is based on a transformer’s encoder ^27^, with additional distance information from the structure module injected at the multi-head attention module in the encoder. The input to the network is first fed into the structure module, and then the pairwise distance matrix of the first 175 amino acids of the α chain and peptide amino acids is calculated using Cβ-Cβ distance. We mask all pairwise distances that are greater than 8Å with a large value. In parallel, we add positional encoding to the input and feed it to the encoder layers together with the pairwise distance matrix. In the multi-head attention module, after calculating the attention logits, we subtract the pairwise distance matrix multiplied by a learnable scalar (Fig. 3C). The masking of all pairs with distance greater than 8Å means they are zeroed in the softmax activation.

### Training and loss functions

The training was performed using ADAM optimizer^41^, with a learning rate of 0.001. The loss function for the structure module was an MSE loss between the predicted and the experimental coordinates after alignment to the reference structure. For the specificity prediction module, the loss function was defined as a binary cross entropy.

## Supplementary data

**Figure S1.**
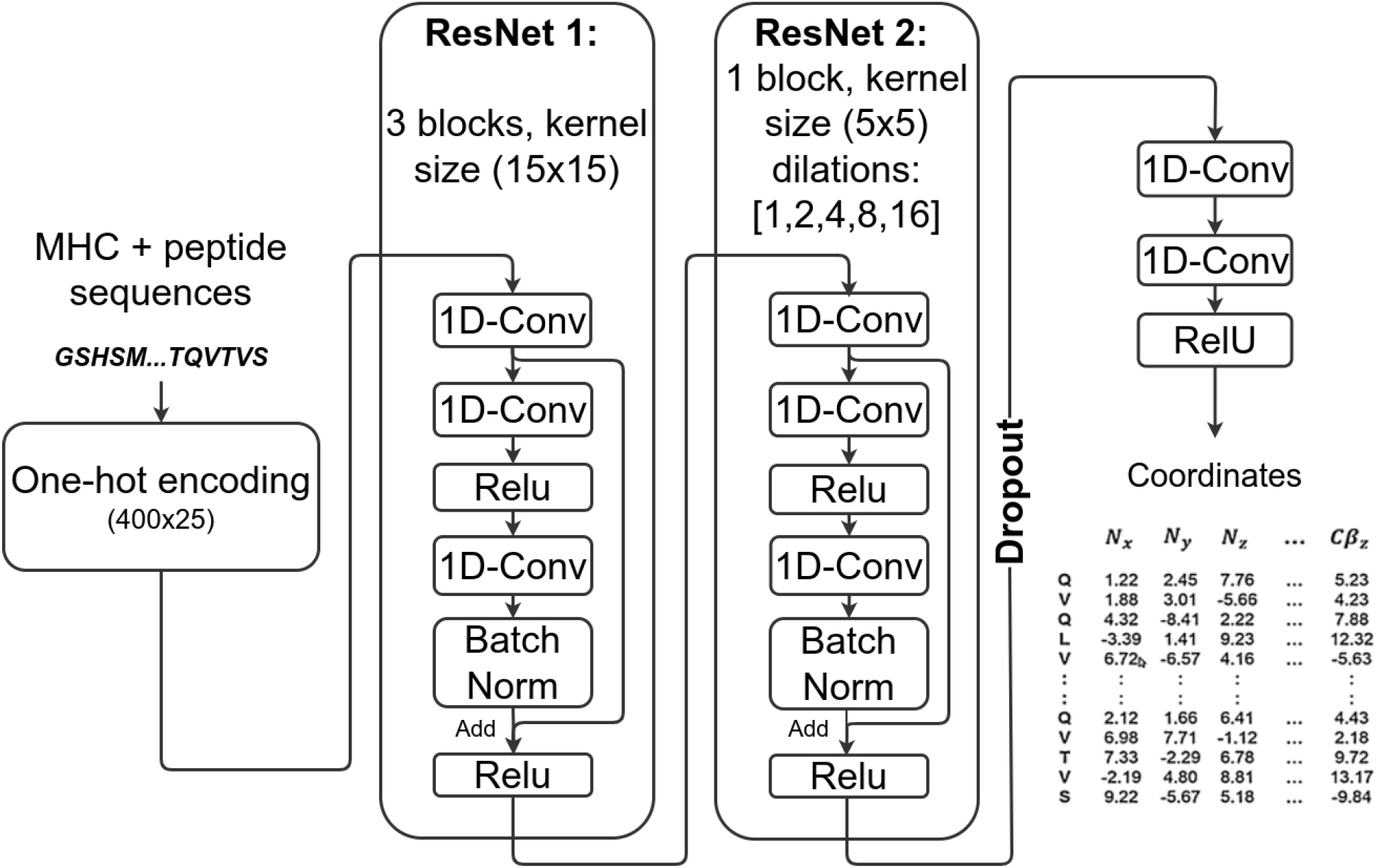
Detailed structure module architecture.

